# Different Paradigms from Computer Vision Align with Human Assessment of the Mouse Grimace Scale

**DOI:** 10.64898/2026.03.04.709492

**Authors:** Michelle Reimann, Jacer Aloui, Niklas Obländer, Niek Andresen, Katharina Hohlbaum, Olaf Hellwich, Patrik Reiske

## Abstract

Animal welfare is a central aspect in animal-based research where mice are most commonly used. Their facial expression can be analyzed to assess their well-being status using the Mouse Grimace Scale. However, its manual application becomes increasingly impractical when used on a large number of animals. This lead to the ongoing integration of computer vision methods to automate the analysis. While such methods have proven effective qualitatively, a systematic assessment to verify their reliability largely remains an open research gap. In this work, we attempted to close this gap as we evaluated three dominant paradigms (i.e., classification from supervised learning features, self-supervised learning features, or landmark locations) for the binary (i.e., well-being un-/impaired) classification of facial mouse images. Our quantitative results showed that such methods can be employed successfully with as low as 16% type II error rates. For qualitative assessment, we visualized the decision-making process and demonstrated that mainly pixels associated with the mouse rather than its environment are used. We further discovered that visual characteristics of the mice beyond those described by the Mouse Grimace Scale contributed to the classification. Our work showed that the automated well-being status assessment in mice is trustworthy and urges towards widespread adoption.

## Introduction

In accordance with ethical principles and good scientific practice, animal-based research requires close monitoring of laboratory animals involved in a study to identify pain, suffering, and distress at the earliest possible point in time. In the European Union, for instance, animals used for scientific purposes are protected by the directive 2010/63/EU that implements the 3R-principle by Russel and Burch^1^ to replace, reduce, and refine the use of animals in research. However, assessing the well-being of laboratory mice (the most widely used animals in research) is challenging as they are nocturnal, show their highest activity at the start and end of the dark period, and usually rest during the light phase^2^. Moreover, it is widely believed that presence of humans may influence their behavior, potentially leading to impaired well-being not being detected^3^. This may be one of the reasons why the emerging trend of continuous monitoring of animals in their home cages using increasingly more automated systems can be observed^4^. For the assessment of mouse well-being, various variables related to behavior, physiology, and external appearance may be surveyed^5,6^. The Mouse Grimace Scale (MGS) can be considered as a behavioral indicator for the well-being of mice. Langford et al.^7^ developed it to assess acute pain from the facial expression of mice: if a mouse is in pain, it will close its eyes and rotate its ears out- or backward; additionally, bulging of the bridge of the nose and cheeks can become visible, and the whiskers may be straightened and drawn back- or forward. These changes in facial expression can be categorized into the five facial action units (FAUs) orbital tightening, nose bulge, cheek bulge, ear position, and whisker change. Once each FAU is evaluated on a discrete three-point scale as absent (score 0), moderately present (score 1) or obviously present (score 2), the overall MGS score is calculated as their mean. In general, an increase in the overall MGS score indicates greater severity of burden. Although the MGS was developed to assess acute pain, some of the FAU can also be altered in other situations, such as after anesthesia, during social interaction or when experiencing fear^8–10^.

The application of the MGS for assessing the well-being of mice is still limited as methods for its automated assessment are scarce and applications for the use in home cage systems have not yet been established. However, such a system would provide multiple benefits, including decreased labor for manual MGS scoring and improved data reliability by mitigating human bias. In addition, continuous 24-hour (real-time) monitoring using standardized methods would enable the early detection of mice with impaired well-being, even at night. First steps towards developing such applications have been made in recent years.

Tuttle et al.^11^ were the first to propose a method for automated pain face assessment in mice. They used an Inception v3^12^ architecture pretrained on the ImageNet^13^ dataset and transfer learned it to solve the binary (i.e., well-being un-/impaired) classification task, that they achieve 94% accuracy on. However, this performance was obtained only after excluding images with low-confidence classifications. Moreover, they only used data showing albino CD-1 mice and no explainability analysis was reported.

Andresen et al.^14^ extended the automated facial expression analysis to black-furred C57BL/6J mice. They compared ResNet-50^15^ and Inception v3^12^ architectures that have been pretrained on the ImageNet^13^ dataset and transfer learned to solve the binary (i.e., well-being un-/impaired) classification task, much like Tuttle et al._11_ did before. Their method achieved an accuracy of up to 99% for detection of post-anesthetic effects caused by injection anesthesia, though cross-treatment generalization was reported as less accurate. They further used deep Taylor decomposition for feature importance visualization, confirming that their model focused on ears, eyes, nose, whiskers, and body features including body posture or piloerection.

Chiang et al.^16^ proposed to use five ResNet-18^15^ models to score the different FAUs separately. Their method achieved 70–88% accuracy in scoring the FAUs individually. However, the reported binary (i.e., well-being un-/impaired) classification accuracy of 63% was lower than that reported in other work. Nevertheless, it should be emphasized that an accuracy of 63% is exactly the same as that achieved by the ground truth (MGS scores assigned by a human scorer with three years of experience) on the same classification task. Much like Andresen et al.^14^ did before, the authors were able to successfully verify that models focus on regions that correspond to appropriate FAUs using Grad-CAM.

McCoy et al.^17^ created PainFace, a cloud-based platform that locates FAUs in a first step to then extract features from these image regions and assess the MGS separately. Their custom multi-branch convolutional network architecture for scoring comprises a total of 18 layers, with the first 12 layers organized into three parallel branches. Using a MGS adopted for black-furred mice, which excluded cheek bulge and scored nose bulge only as absent (score 0) or clearly present (score 2), they reported 84–99% precision for bounding box prediction and an F_1_ score of 0.75–0.84 for pain classification depending on the FAU. The accuracy was not reported. Different from other work, McCoy et al.^17^ did not report any systematic explainability validation.

Gupta et al.^18^ proposed an alternative quantitative approach for pain detection replacing the MGS. They used DeepLabCut (DLC)^19^ to extract landmark locations on the right eye of Nav1.8ChR2 mice and to calculate the distance between the eyelid margins and palpebral fissure width. Excluding images in which any landmark had a likelihood below 0.95, they reported a significant reduction of the average eyelid distance in the frequency distribution across different treatments, as well as a smaller but still significant reduction in the average palpebral fissure width for most treatments.

Kobayashi et al.^20^ proposed to learn a model for pain face assessment under weak supervision only using metadata for labels, assuming that well-being is unimpaired pre-surgery and impaired post-surgery. They used a custom model architecture that may be considered a simplification of the ResNet-18^15^ architecture, as it follows the same general structure but is not as deep nor uses all the same components. They reported that their method achieved 97.0% in sensitivity and 99.4% in specificity on the training data, and that it successfully generalizes to capsaicin- and CGRP-induced pain models not seen during training. However, they further reported that their model did not generalize to black-furred C57BL/6 mice, suggesting that strain-specific models may be required. Here, it is important to note that reporting performance on training data may impede reliable assumptions about the performance on previously unseen data. Much like Chiang et al.^16^ before, the authors used Grad-CAM to successfully verify that the learned models base the MGS estimation on the FAU regions.

Sturman et al.^21^ proposed GrimACE, the first end-to-end hardware-software system for standardized MGS assessment combining automated image acquisition, scoring, and full-body pose estimation. Using a Vision Transformer^22^ model pretrained on the ImageNet^13^ dataset and fine-tuned on human-annotated images, it achieves high correlation (Pearson’s r=0.87) with expert human scorers. Crucially, by standardizing image acquisition, GrimACE addresses a fundamental limitation of prior work: without controlled imaging conditions, training data for different treatments is often acquired across different laboratories, introducing hidden biases that compromise treatment comparisons. The system was validated in a post-surgical craniotomy model, successfully detecting moderate pain levels. However, no explainability analysis of the model’s decision-making process was reported.

Most recently, Andresen et al.^23^ provided a large, diverse dataset of mouse face images varying in terms of mouse strain, experimental treatment, animal facility, and image acquisition setup in a follow-up study to their earlier work. This dataset was used to train a ResNet-50^15^ model to predict the mean MGS score as a floating point number between 0 and 2. Depending on the subsets of this diverse dataset included in the model training, a root mean squared error of up to 0.26 was reached and high correlation (Pearson’s r=0.85) between the human scorers and the MGS scores that were automatically predicted was achieved. Andresen et al.^23^ demonstrated that a limitation to the orbital tightening FAU did not improve model performance; instead, it resulted in a deterioration.

While these methods demonstrated that automated MGS assessment is feasible, the available literature lacks systematic comparison across paradigms under standardized conditions as each of them employs different datasets, preprocessing pipelines, and evaluation protocols, precluding direct assessment of how methodological choices affect both predictive performance and model interpretability. In this work, we addressed this gap by comparing three computer vision paradigms on a large, diverse dataset of mouse face images including MGS scores: classification using supervised learning features directly from facial images, the dominant approach in prior work; classification from anatomical landmark locations, leveraging pose estimation frameworks that are increasingly standard in behavioral research groups; and classification from self-supervised representation learning features, which learns visual features from images without human annotations. These three dominant paradigms were trained on a binary (i.e., well-being un-/impaired) classification task to predict whether the well-being of a mouse shown in an image was impaired or unimpaired. Crucially, we complement quantitative evaluation with systematic explainability analysis, examining whether models trained following different paradigms converge on biologically meaningful facial features.

## Results and discussion

### Quantitative performance analysis

To quantitatively assess model performance the binary (i.e., well-being un-/impaired) classification task, we learned the parameters in five independent runs per paradigm, as described in the methods section. In Tab. 1, we reported macro-averaged precision (i.e., positive predictive value; the number of correctly classified images divided by the number of images correctly or incorrectly classified as a class, averaged over all classes), recall (i.e., sensitivity; the number of correctly classified images divided by the total number of images of a class, averaged over all classes), and F_1_ score (i.e., harmonic mean of precision and recall; two times the product of precision and recall, divided by their sum) on the random holdout test split of our dataset for each of the three tested paradigms (i.e., classification from supervised learning features, self-supervised learning features, or landmark locations) separately. The F_1_ score is useful when a balance between precision and recall is needed. All three of these metrics take values from zero, indicating the lowest performance, to one, indicating the best performance. As can be seen there, the classification from self-supervised learning features reached the highest mean precision, recall, and F_1_ score at 0.83, 0.84, and 0.83, respectively. The classification from supervised learning reached almost as good performance at 0.80 for all three metrics, while the classification from landmark locations achieved the lowest performance at 0.68 precision, 0.68 recall, and 0.63 F_1_ score. The low standard deviation of 0.01 to 0.02 in all metrics suggests that the reported metrics reliably reflect the performance of the three tested paradigms.

**Table 1.** Precision, recall, and F_1_ score (mean±std) macro-averaged over five runs on holdout data, where bold font marks the best performance per metric.

| Classification from | Precision | Recall | F<sub>1</sub> score |
| --- | --- | --- | --- |
| Supervised learning features | 0.80±0.01 | 0.80±0.01 | 0.80±0.01 |
| Self-supervised learning features | **0.83±0.01** | **0.84±0.01** | **0.83±0.01** |
| Landmark locations | 0.68±0.02 | 0.68±0.01 | 0.63±0.02 |

To assess the performance of the three different paradigms in more detail, Tab. 2 reports their precision, recall, and F_1_ score separated by binary (i.e., well-being un-/impaired) class. The classification from self-supervised learning features again reached the highest precision, recall, and F_1_ score values for both classes, “well-being impaired” and “well-being unimpaired.” Also again, the classification from supervised learning features achieved almost as high precision, recall, and F_1_ score values while the classification from landmark locations fell behind. Here it is important to note that for all paradigms the performance on the “well-being unimpaired” was higher than for the “well-being impaired” class. However, in context of animal experiments the “well-being impaired” class is more important to be scored correctly since animals experiencing pain, suffering and/or distress should be identified as quickly as possible to treat them appropriately. We assessed the type II error (i.e., false negative rate; one minus recall) for the “well-being impaired” class for either of the three tested paradigms. The type II error is the probability that an image showing a mouse with “impaired well-being” is incorrectly classified as “well-being unimpaired.” The classification from self-supervised learning reached the lowest type II error at 0.16, with the classification from supervised learning features only reaching a slightly higher rate of 0.20, and the classification from landmark locations falling behind at a much higher 0.36 false negative rate. Again, the low standard deviation of 0.01 to 0.04 in all metrics suggested that the reported metrics reliably reflect the performance of the three tested paradigms.

**Table 2.**
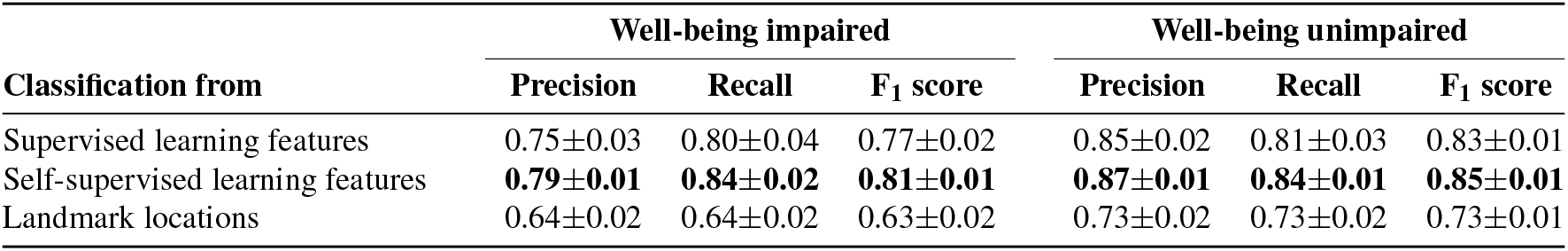
Precision, recall, and F_1_ score (mean±std) averaged per class (“well-being impaired”/”well-being unimpaired”) over five runs on holdout data, where bold font marks the best performance per class per metric.

| Classification from | Well-being impaired |  |  | Well-being unimpaired |  |  |
| --- | --- | --- | --- | --- | --- | --- |
|  | Precision | Recall | F<sub>1</sub> score | Precision | Recall | F<sub>1</sub> score |
| Supervised learning features | 0.75±0.03 | 0.80±0.04 | 0.77±0.02 | 0.85±0.02 | 0.81±0.03 | 0.83±0.01 |
| Self-supervised learning features | **0.79±0.01** | **0.84±0.02** | **0.81±0.01** | **0.87±0.01** | **0.84±0.01** | **0.85±0.01** |
| Landmark locations | 0.64±0.02 | 0.64±0.02 | 0.63±0.02 | 0.73±0.02 | 0.73±0.02 | 0.73±0.01 |

For an automated system, it is crucial that the type II error is as low as possible. If an automated system indicates that the well-being of a mouse is reduced, a responsible person (i.e., the Laboratory Animal Science-staff, veterinarian or researcher) is required to perform a health check to confirm the well-being status of a mouse and, in a case of compromised well-being, ensure that appropriate measures are taken to alleviate its burden. No measures need to be taken if its well-being is not disturbed; false positive cases (i.e., mice whose well-being is not reduced are classified as “well-being impaired”) may increase the workload and potentially cause mild stress to the animals because of the health check. However, the consequences of false negative cases (i.e., mice whose well-being is reduced are classified as “well-being unimpaired”) are much more serious for animal welfare when using an automated system because the burden of these animals remain undetected, unless additional checks are performed by a responsible person. Therefore, a low type II error should be aimed for.

### Qualitative assessment of model attention

For qualitative assessment, we employed layer-wise relevance propagation^24^ as Otsuki et al.^25^ proposed. The output of the model is propagated through the network in reverse order, attributing relevance scores to input pixels of an image. The relevance scores are displayed as heatmaps that highlight the visual features in an image that were used in the decision-making process (i.e., to classify an image as “well-being impaired” or as “well-being unimpaired”). This method was utilized for both the supervised learning model and the self-supervised learning model. Due to the low performance of the landmark location based classification, which was demonstrated in the quantitative performance analysis and which we deem too low for any real-world application, it was excluded from the qualitative assessment of model attention discussed in this section.

The visualization demonstrated that the decision of the models was based primarily on the mouse rather than its environment. Similar findings regarding the feature importance visualization of models trained on this particular binary (i.e., well-begin un-/impaired) task were previously reported^14^. In the face, characteristics known from the MGS^7^ in combination with additional features were identified: the eyes and fur around the eyes, the ears (i.e., parts of their outline, hair around/in front of the ears, opening of the external auditory canal), the whiskers and the whisker pad, the nose tip, and the ventral outline of the snout/upper lip.

Orbital tightening is one of the FAUs of the MGS, as mice generally close their eyelids when their well-being in impaired. In addition to the degree of eye closure, the fur around the eyes may be relevant (Fig. 1a), since an eye squeeze causes wrinkles around the eyes. Another explanation for the importance of the area around the eye may be the use of eye drops or cream during anesthesia/surgery, which makes the fur appear wet and slightly disordered. The outline of the ears (Fig. 1b) forms lines and angles which, possibly in combination with the outline of the body and/or head, may allow conclusions to be drawn about the position of the ears and their distance. Furthermore, these lines and angles may indicate whether the ears are rounded and oriented forward, or whether they appear more folded and pointed. These changes in ear position are also partially described in the MGS. In images taken from the side view, the opening of the external auditory canal is visible when the ears are rotated outwards (Fig. 1c) and seemed to play a critical role for the models. It is conceivable that the models only take into account the fact that the opening of the external auditory canal is visible from the side, or even the angles of the lines and the visible area. For instance, a smaller area is visible when the ears are folded, indicating impaired well-being. Moreover, both the whiskers and the whisker pad contributed to the decision-making process (Fig. 1d). The appearance of the whisker pad may change when the muscles connected to the vibrissae follicles contract^26^, causing the whiskers to straighten and draw backward or forward, as observed in mice with impaired well-being and described in the MGS. Another feature of interest was the tip of the nose (Fig. 1e), especially for images assigned to the “well-being unimpaired” class. As already considered by Andresen et al.^14^, its color and position may be associated with a different well-being state. After anesthesia, surgery, or the application of certain substances or in cold stress, the tip of the nose may turn paler and point downwards due to the hunched body position^14^. In contrast, a mouse with undisturbed well-being is more explorative and, therefore, the chance that the nose is raised in the air in an image is higher.

**Figure 1.**
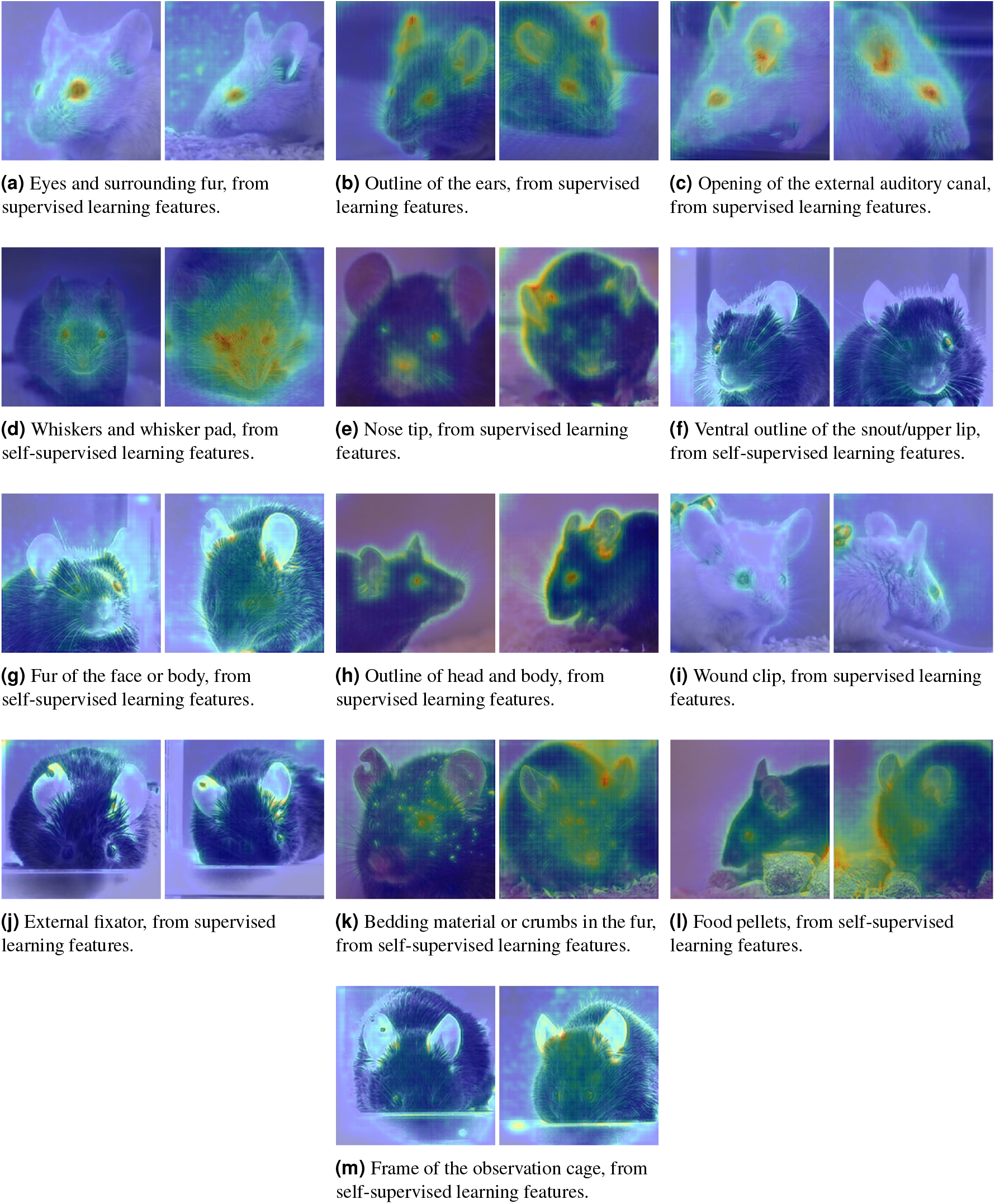
Layer-wise relevance propagation: example pairs of images correctly classified as “well-being unimpaired” (each left) or “well-being impaired” (each right) side-by-side for the models paying no to little (blue) or more (from yellow to red) attention to different visual cues.

A novel facial feature identified by the visualization was the ventral outline of the snout/upper lip (Fig. 1f). If the snout/upper lip has a rather round shape, it seems to be relaxed and indicate unimpaired well-being. In contrast, impaired well-being is more associated with a tense snout/upper lip, resulting in a more angular shape.

The self-supervised learning model appeared to also use the fur of the face and the body (Fig. 1g): if the well-being of a mouse is impaired, the mice may show piloerection (i.e., the muscles at the base of the hair follicles contract, causing the hair to stand up)^27^. Moreover, it is conceivable that the models detected the bulges on the bridge of the nose and the cheeks caused by the experimental procedures, indicating impaired well-being. Another noteworthy observation was that the self-supervised learning model primarily focused on features visible in the foreground of the image, whereas the supervised learning model concentrated on features located in the background. For instance, in an image captured from the animal’s left side, the self-supervised learning model mainly incorporated pixels from the left ear into its decision-making process and the supervised learning model predominantly relied on pixels from the right ear.

For some images, it appeared that not the entire coat of the mouse but only the outline of body and/or head was incorporated into the decision-making process (Fig. 1h). If the well-being of a mouse is impaired, it may show a hunched body posture^27^ and the head tends to drop downwards, with the tip of the nose pointing downwards. This observation suggests that the models may take into account these lines revealing the body and head position, as discussed previously by Andresen et al.^14^.

Beyond these features relating to the mouse, however, the models also relied on other visible elements that may be associated with the well-being status of the animal. In the subset JW, the surgical wound was closed with a wound clip (Fig. 1i), and in the subset AW, an external fixator was attached to the left femur (Fig. 1j), indicating that surgery had been performed previously. In contrast to the subset JW and AW, no surgical fixation devices were visible in images of the subset KH. Images of this subset were recorded in an observation cage with bedding scattered on the floor. Additionally, the mice had access to a tube filled with food pellets, encouraging mice to burrow. After burrowing, bedding material or crumbs of food pellets were found in the faces of the animals (Fig. 1k). Mice whose well-being is impaired display reduced or no burrowing behavior^28^. Therefore, bedding material or crumbs visible in the mice’s faces may suggest that the well-being of the animal is not impaired. Interestingly, the models included these features that may point to a post-anesthetic or post-sugrical period in the decision-making process for both classes; thus the wound clip, the external fixator or bedding material/crumbs in the faces may have not primarily contributed to the distinction between the two classes “well-being impaired” and “well-being unimpaired.” In rare cases for the subset KH, the models used the food pellets the mice removed from the tube (Fig. 1l). However, this occurred with both classes. In the subset AW, the frame of the observation cage was visible in several images and was highlighted in the heatmaps of images assigned to both classes (Fig. 1m). Considering that the majority of the model parameters were not learned on image data showing mice in laboratory settings specifically, we assume that such visual cues are being paid attention to because they are rather rare outliers. Notably, the models do not learn to overfit either binary (i.e., well-being un-/impaired) class to the presence of such a signal.

Attention maps for all images used in the present study are publicly available at https://doi.org/10.14279/depositonce-25407 and the original images at https://doi.org/10.14279/depositonce-25192.

## Conclusion

We used the three dominant computer vision paradigms (i.e., classification from supervised learning features, classification from self-supervised learning features, and classification from landmark locations) to recognize whether the well-being of mice is impaired or unimpaired from their faces. For this binary (i.e., well-being un-/impaired) classification, a diverse dataset containing images from different mouse strains, treatments, and image acquisition setups was utilized to simulate the diversity of experimental or housing conditions in a real-world application. Classification from self-supervised learning features and supervised learning features achieved a high performance, and outperformed classification from landmark locations. Both the classification from self-supervised learning features and the classification from supervised learning features mainly used pixels associated with the mouse rather than its environment for the decision-making process: facial features known from the MGS and additional visual characteristics of the mice contributed to the decision of the paradigms. Since humans with experiences in severity assessment in mice also rely on these visual characteristics, the performance of the paradigms has proven to be reliable. For the application of the proposed semi-automated pipeline, only minimal computational resources are required, making it accessible to researchers in the field of animal-based research and encouraging them to adopt and further develop this approach.

Our interdisciplinary approach, bridging computer vision and veterinary medicine, enabled us to validate the reliability of the paradigms and to identify novel characteristics for well-being assessment, such as the ventral outline of the snout/upper lip, that have not been previously described and warrant further investigation.

## Methods

### Animals

For the present study, no additional animal experiments were conducted. Instead, we reused the dataset of mouse face images that Andresen et al.^23,29^ published previously. All relevant information on the licensing committee approving the animal experiments, permit numbers, guidelines and regulations, as well as information required by the ARRIVE guidelines^30^ are provided in their work as well as the underlying studies^8,9,31–36^ in which the animal experiments were carried out.

### Dataset

The dataset used in this work was presented by Andresen et al.^23,29^ and consists of almost 35,000 facial images of mice. The images were collected from five research groups (AW, JW, KH, LW, and MR) conducting independent studies and show five mouse strains (BALB/c, C57BL/6J, C57BL/6N, NMRI, and DBA1) of both sexes. The treatments (anesthesia, surgery, injections of substances, experimental housing conditions) varied in severity. The dataset is not balanced: laboratories, mouse strains and sexes, treatments, and facial expressions are represented unevenly. For about 3,400 images, the dataset provides human-generated MGS scores annotated by one to five human experts each. Besides discrete MGS scores 0, 1, and 2 (i.e., signs of impaired well-being not, moderately, and obviously present) per FAU, these labels also indicate whether the human expert was unable to score a FAU, possibly due to blurring or occlusion in the image.

#### Data cleaning

We only considered labeled images, as our work is set to evaluate three computer vision paradigms using models that have been pretrained to process images into features used for classification. Our protocol for learning the classifiers is reported in this work’s methods section, their evaluation in its results and discussion section.

In earlier work, Hohlbaum et al.^37^ reported that of all FAUs orbital tightening is most reproducible (i.e., has the highest agreement between human scorers) and Ernst et al.^38^ that orbital tightening is the most reliable FAU to assess for examining pain induced by intraperitoneal CCl_4_ injections. Therefore, we cleaned the dataset from images where less than three of five FAUs are visible (i.e., have been assigned human-annotated scores) and make the visibility of the orbital tightening FAU a necessary condition in order to investigate whether or not different computer vision paradigms align with human assessment.

The resulting dataset consists of 3,286 labeled images, which is about 10.6% of the full dataset.

#### Calculating binary labels from Mouse Grimace Scale scores

As this work is concerned with binary (i.e., well-being un-/impaired) classification, the mean MGS score was used to assign the binary labels to the images, as follows:

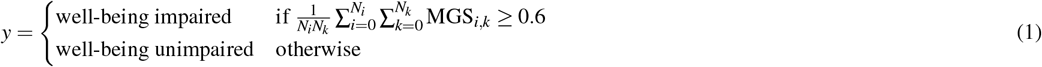

where *N*_*i*_ is the number of human experts that have annotated MGS scores, *N*_*k*_ is the number of valid (i.e., FAU visible and scored) human annotations, and 0.6 is the threshold value for the binary (i.e., well-being un-/impaired) decision.

This process resulted in 1,389 images being labeled as “well-being impaired” and 1,897 as “well-being unimpaired,” making for an almost balanced dataset.

### Computational resource requirements

The computational requirements for any of the tested computer vision paradigms were rather low as all of them made use of pretrained model parameters. To transfer learn the models used in our experiments, we used NVIDIA graphics cards. However, these where primarily used to accelerate the learning process and not for their memory. The actual learning process of the more resource-intensive ResNet-50^15^ models took about 15 minutes and only used just over 3GB of graphics memory. Considering these low computational resource requirements, we consider hardware (e.g., graphics cards) dedicated to learning deep neural networks not strictly necessary. Instead, any reasonably powerful desktop or laptop computer (as are usually already used in any research facility) is powerful enough to learn such a model.

### Classification from supervised learning features

We used a ResNet-50 model that has been pretrained under full supervision^15^ on the ImageNet^13^ dataset as feature encoder, as others (e.g., Tuttle et al.^11^ and Andresen et al.^14^) did before. To that end, the pretrained output layer was replaced by a linear layer with randomly initialized parameters that maps the model’s 2,048 features to 2 output values to solve the binary (i.e., well-being un-/impaired) classification problem. To learn this layer’s parameters, we randomly split the dataset into two disjunct subsets: 70% only used for training and 30% only for testing. We used the cross entropy criterion, weighted to counteract the binary (i.e., well-being un-/impaired) class imbalance in the dataset; the Adam^39^ optimizer with a 0.001 learning rate, betas 0.9 and 0.999, and no weight decay for batches of 128 images each; the learning rate was first increased following a linear warm-up schedule from starting factor 0.001 for 10 epochs, before it was decreased following a cosine annealing schedule for another 90 epochs. Input images were resized such that their short side measured 224 pixels before a center crop of 224×224 pixels was performed. In a final step, color channels were normalized to have (0.485, 0.456, 0.406) means and (0.229, 0.224, 0.225) standard deviations.

### Classification from self-supervised learning features

Much like in case of the classification from supervised learning features, we used a ResNet-50^15^ model that has been pretrained on the ImageNet^13^ dataset under self-supervision using the Barlow Twins^40^ criterion as feature encoder. We used a final linear layer with randomly initialized parameters to solve the binary (i.e., well-being un-/impaired) classification problem by mapping the model’s 2,048 features to 2 output values; randomly split the dataset into two disjunct subsets, 70% only for training and 30% only for testing; weighted the cross entropy criterion to counteract the binary (i.e., well-being un-/impaired) class imbalance in the dataset and optimized it using the Adam^39^ optimizer with a 0.001 learning rate, betas 0.9 and 0.999, and no weight decay with a 128-image batch size and the learning rate following a 10-epoch warm-up schedule from starting factor 0.001 before following a cosine annealing schedule to decrease it over the following 90 epochs. Again, input images were resized to measure 224 pixels on their short side before the center 224*×*224 pixels were cropped, followed by normalizing the color channels to have (0.485, 0.456, 0.406) means and (0.229, 0.224, 0.225) standard deviations.

### Classification from landmark locations

The binary (i.e., well-being un-/impaired) classification of mouse faces from landmark locations requires those to be reliably located in the input images. Despite the available literature not defining a set of landmarks that is commonly used for this task, it has been reported^37,38^ that orbital tightening is the most relevant FAU to assess. This was later also verified by Gupta et al.^18^ for automated MGS assessment. Therefore, we defined the medial and lateral canthus, as well as the halfway point between the two on the upper and lower eyelid for both eyes as landmarks representing orbital tightening. Of the remaining FAUs, only the ear position is rather robust against common challenges (e.g., blurriness) found in image data. We therefore added the apex of auricle, as well as 2 points each on the medial and lateral margin of the auricle as landmarks representing the position for both ears. To enable normalization of the landmark locations, we added the tip of the nose as landmark. Fig. 2a depicts the 19 landmarks used in our work.

**Figure 2.**
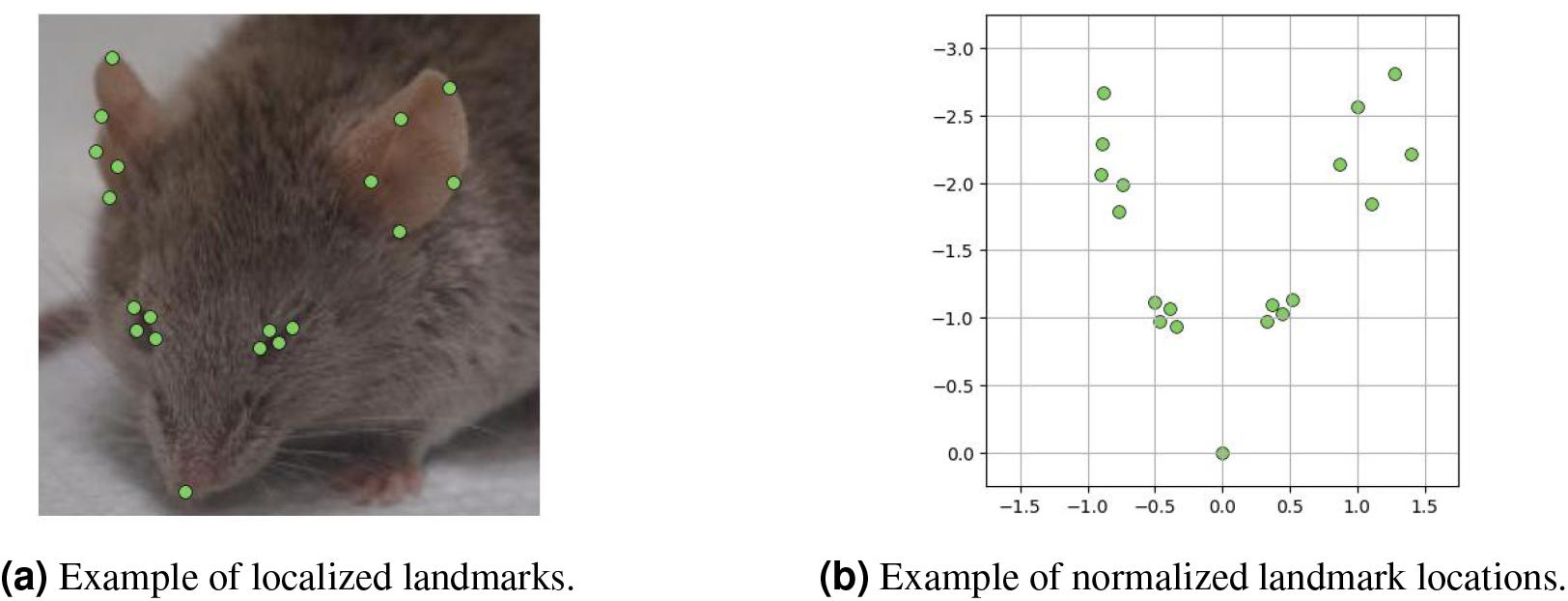
Example for the localization and normalization of landmarks.

For landmark localization, we learned a DLC model by Mathis et al.^19^ on a set of 430 images from our dataset that have manually labeled landmarks. These images were selected to represent the diversity of the dataset: not just with respect to laboratories, mouse strains, etc. but also sampling from different combinations of annotated MGS scores. To increase landmark location accuracy, we used the HRNet-W18^41^ network architecture, which was pretrained on ImageNet^13^ and transfer learned it for a total of 50 epochs on 408 randomly selected images only used for training while testing it on the holdout data. We followed the DLC (version 3.0.0) standard recipe to fine-tune the HRNet-W18 model parameters using the PyTorch^42^ framework. This includes a heatmap head for landmark detection with location refinement. We optimize the weighted mean squared error criterion (weight: 1.0) as heatmap loss and the weighted Huber criterion (weight: 0.05) for location refinement using the AdamW optimizer with a 0.0001 learning rate, and no weight decay for batches of 16 images each. We augmented the images by applying random horizontal flipping, random rescaling by random factors of 0.8 to 1.25, random translation by 5 pixels, random rotations of up to 30 degrees, random Gaussian noise with a 12.75 standard deviation and enabled motion blur with a kernel of size 3 to 7. Cropping was applied on all images with a crop shift of 10% to obtain images of size 224×224, followed by normalizing the color channels to have (0.485, 0.456, 0.406) means and (0.229, 0.224, 0.225) standard deviations.

As the dataset contains mouse facial images from different perspectives, different alignment procedures are required due to occlusions that depend on head orientation. As no standard normalization factor exists for mouse facial landmarks, we applied a similar perspective-dependent strategy as Pessanha et al.^43^ did in their work for horses. In order to determine a non-obstructed reference eye, facial perspective is inferred from landmark visibility (i.e., DLC confidence scores). The right eye was used as the reference by default; if fewer than three landmarks of the right eye exceeded a confidence threshold of 0.5, the left eye was used instead. Upon successful landmark localization and selection of a reference eye, all landmark coordinates were normalized: all coordinates are translated such that the nose tip lies in the coordinate origin; rotated such that the nose–eye line lies at an angle of 70 or 110 degree for the right-hand or left-hand side of the face, respectively; and scaled such that the nose-eye line is of unit distance. Fig. 2b depicts this normalization.

The resulting normalized coordinates were then used as input for linear probing to perform binary (i.e., well-being un-/impaired) classification. Each of the 19 facial landmarks was represented as a tuple of two-dimensional (x and y) coordinates and input to the classifier as a vector of 38 dimensions. Again, we randomly split the dataset into two disjunct subsets, 70% only for training and 30% only for testing; weighted the cross entropy criterion to counteract the binary (i.e., well-being un-/impaired) class imbalance in the dataset and optimized it using the Adam^39^ optimizer with a 0.001 learning rate, betas 0.9 and 0.999, and no weight decay with a batch size of 128 images, and the learning rate following a 10-epoch warm-up schedule from 0.001 starting factor before following a cosine annealing schedule to decrease it over the following 90 epochs.

In an ablation study, our landmark locations based experiments were also conducted using a nonlinear multi-layer network architecture similar to that Feighelstein et al.^44^ used. But despite its potential to learn a more complex task than a linear layer (i.e., linear probing) alone, its performance remained inferior. Unfortunately, Feighelstein et al.^44^ did not conduct an ablation study comparing their architecture against linear probing. However, we notice that we utilized fewer, less redundant landmarks than they did. For instance, they used a total of eight two-dimensional landmarks per eye and pupil whereas we only used four per eye and ignore the pupil. We therefore hypothesized that nonlinear architectures as theirs are only beneficial if the input signals carry highly redundant information such that dimensionality reduction is required. Due to its inferior performance, we omitted experiments using a nonlinear multi-layer classifier^44^.

## Acknowledgements

We thank Yiwei Diao, Patrick Freund, Qinyi Hui, Anabel F. Kasprick, Florentin Marquardt, and Mirco Richter for their contribution to this work in its early stages; we thank Lena Reiske for critically proofreading our manuscript before submission.

This research was funded by the Deutsche Forschungsgemeinschaft (DFG; lit. German Research Foundation) under Germany’s Excellence Strategy – EXC 2002/1 “Science of Intelligence” – project number 390523135.

## Author contributions

P.R. conceptualized this work; M.R., J.A., N.O., and P.R. conceived and conducted the experiments; N.A., K.H., and P.R. analysed the results; M.R., J.A, and N.O. drafted the manuscript; N.A., K.H., O.H., and P.R. edited the manuscript.

## Data availability

The data used in this research was presented by Andresen et al.^23,29^ and is publicly available at https://doi.org/10.14279/depositonce-25192 while the data we produced in our qualitative experiments will be made publicly available at https://doi.org/10.14279/depositonce-25407 upon publication. Our training logs and learned model parameters will be made available upon reasonable request.

## Code availability

Our source code will be made available upon reasonable request.

